# Catalytic rate constant for the utilization of biopolymers

**DOI:** 10.64898/2026.05.29.728646

**Authors:** Ikechukwu Iloh Udema

**Affiliations:** Department of Chemistry and Biochemistry, Research Division, Ude International Concepts LTD (862217), B. B. Agbor, Delta State, Nigeria

**Keywords:** Human salivary alpha amylase, Invertase, First-order rate constant for product production, First-order rate constant for the utilization of substrate, Turnover number

## Abstract

The catalytic rate constant (*k*_*cat*_) for product formation is considered a turnover number. Therefore, it is often mistakenly believed that *k*_*cat*_ equals the turnover number and the number of substrate molecules changed per unit of time. Therefore, the aim of this study is to show that the rate constant for product synthesis and release is not always the same as the rate constant 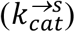 for substrate utilization. To determine the precise substrate concentration at which these two rate constants are identical, it is appropriate to derive equations that allow the computation of 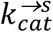. In the end, the study will provide the most likely concentration of enzymes that can guarantee minimal or no recycling. An analysis of the literature on invertase (EC 3.2.1.26) and the Bernfeld method of generating Michaelian kinetic parameters for human salivary alpha-amylase (HSAA, EC 3.2.1.1) revealed that all kinetic parameters except 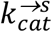 increased with substrate concentration. Meanwhile, the values for invertase decreased from 0.0697 to 0.0361/min, and the values for HSAA decreased from 5,802.4687 to 3,213.0124/min. The magnitude of 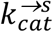 for each substrate concentration ([*S*_*T*_]) is not always equal, except when [*S*_*T*_] is determined post-assay by computation or extrapolation. The lower [*S*_*T*_] at which 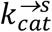 and *k*_*cat*_ for [HSAA] are equal is 3.667540128 g/L (5.682584642 M), which is similar to the molarity of HSAA (5.6101967709 M). The *k*_*cat*_ for HSAA was 11,930.9885/min. Future assays should aim to generate large amounts of data for a robust statistical analysis.

**Graphical abstract:** 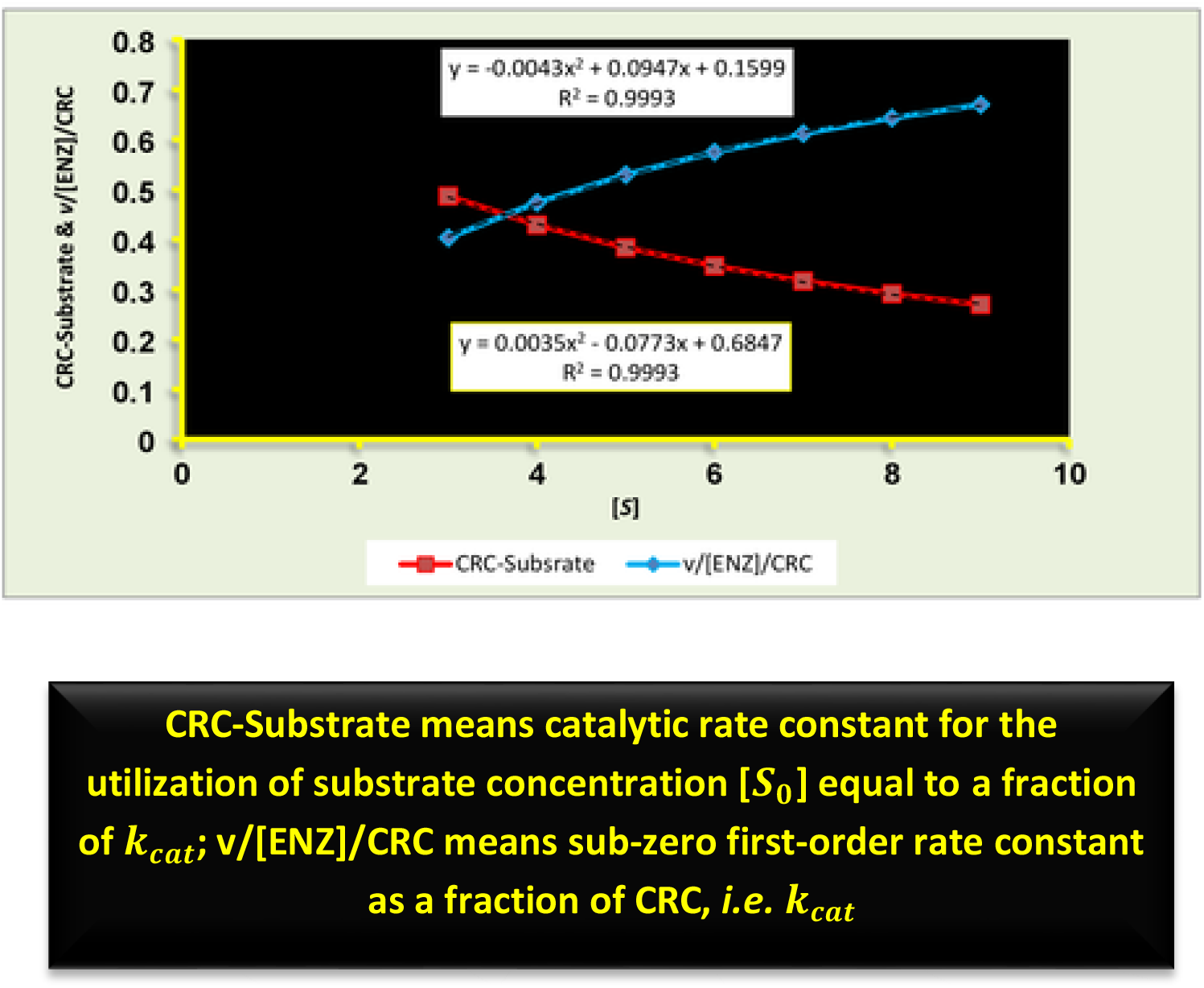

## 1. Introduction

In order to understand the mechanics behind various enzyme-catalyzed reactions, studies in the literature have looked at turnover and turnover rates. The turnover number and the number of substrate molecules transformed per unit of time are frequently thought to be equal to the catalytic rate constant (*k*_*cat*_) for product synthesis. Some works investigate single-turnover dynamics without explicitly specifying *k*_*cat*_ values, despite criticism of their use. If the latter is stated, it creates the impression that it is a typical Michaelian zero-order kinetic constant, as opposed to a sub-Michaelis-Menten kinetic constant, which is much smaller. The works of Bergman and Sousa (2025) are a prime example of this. Using a fluorescent reporter of OMP folding (*i*.*e*., bOmpA-A488), single-turnover kinetic parameters were generated for the wild-type BAM complex *in vitro*: a *k*_*fold*_ of (0.78 ± 0.15)/min and an approximate substrate affinity of 3.1 ± 1.1 μM (Bergman & Sousa, 2025). These values are sub-zero-order kinetic parameters, *i*.*e*., they are not Michaelian. Rather, *k*_*fold*_ is less than zero-order kcat, while the substrate affinity, also referred to as the Michaelis–Menten constant, is actually more of an enzyme–substrate dissociation constant. This is because a single-turnover assay demands that the enzyme concentration be several times greater than the substrate concentration.

Notable of some of such include studies on conformational transitions during the interactions of AlkB with the ds15m^1^A substrate *via* Trp and 2aPu fluorescence and FRET by means of single turnover kinetics of dsDNA demethylation by AIkB. Also, conformational dynamics of the AlkB protein during interactions with the ss15m^1^A substrate has been studied *via* Trp fluorescence under single-turnover conditions (Kanazhevskaya *et al*., 2019). Conformational transitions during the interactions of AlkB with the ds15m^1^A substrate has been studied *via* Trp and 2aPu fluorescence and FRET by means of single turnover kinetics of dsDNA demethylation by AIkB. Also, conformational dynamics of the AlkB protein during interactions with the ss15m^1^A substrate has been studied *via* Trp fluorescence under single-turnover conditions (Kanazhevskaya *et al*., 2019).

Based on turnover kinetics, the kinetic mechanisms of class A and class C β-lactamase catalysis have been studied. The results indicate that the mechanistic differences can be explained by the way these enzymes catalyze the hydrolysis of penicillins and cephalosporins (Adediran & Pratt, 2022). Although the term “turnover” refers to the number of substrate molecules transformed per unit time in most of the literature, it may also imply the catalytic rate constant for product production and release. A kinetic and mechanistic framework for R2 non-LTR Retrotransposition has been studied based on a single turnover kinetic analysis with global data fitting which defined the rate constants for each step in the pathway (Dangerfield *et al*., 2026). The development of a single turnover stopped-flow fluorescence strategy that reports on processive protein unfolding catalyzed by caseinolytic peptidase B (ClpB) have been reported (Banwait *et al*, 2024).

So, a key aim of this study is to show that the rate constant for product synthesis and release is not always the same as the rate constant for substrate usage. To determine the precise substrate concentration at which these two rate constants are identical, it is appropriate to derive equations that allow the computation of the substrate utilization rate constant 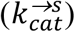, which is different from *k*_*cat*_. In the end, the study will provide the most likely concentration of enzymes that can guarantee minimal or no recycling.

## 2. Theoretical development

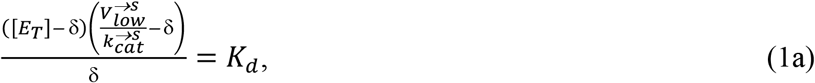

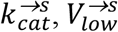, [*E*_*T*_], δ, and *K*_*d*_ are the catalytic rate constant for the transformation of the substrate (S) to product (P), maximum velocity of the transformation of S to P, molar concentration of the enzyme (E), molar concentration of the substrate that binds the same concentration of E, and the equilibrium dissociation constant respectively. Realize that Eq. (1a) arises on the premise that 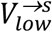 is directly proportional to [*S*_0_]/*M*_*s*_ at the first 3 to 4 different concentrations of the substrate within the range of substrate concentrations explored for the assay. Thus, 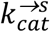 is equal to 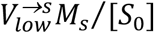. First rearrangement of Eq. (1) gives the following:

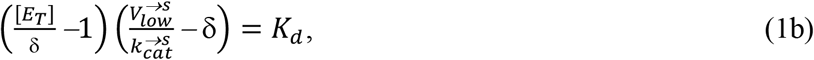

Expansion gives the following:

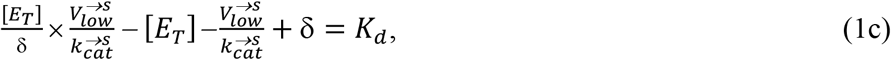

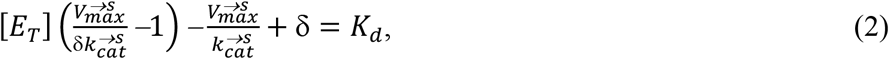

If δ = *v*/*k*_*cat*_, then,

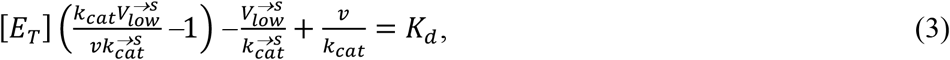

Solving for [*E*_*T*_] gives the following:

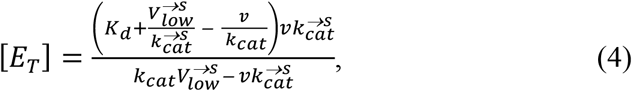

Next step is to generate Lineweaver-Burk equation from the equation as follows:

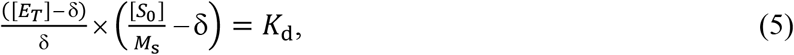

Where [*S*_0_] and *M*_s_ are the mass concentration of the substrate and its molar mass respectively. Expansion of Eq. (5) gives:

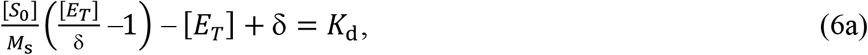

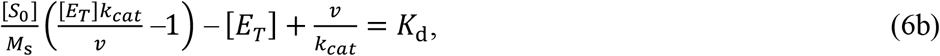

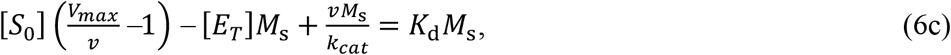

Solving for [*E*_*T*_]gives the following:

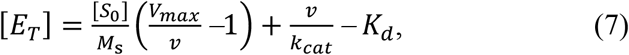

Thus, Eqs (4) and (7) are the same. Hence,

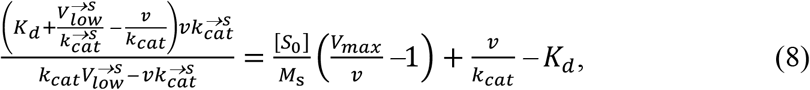

Expansion of Eq. (8) gives the following:

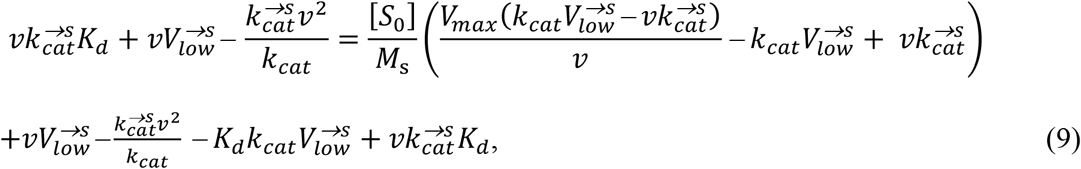

Simplifying gives the following:

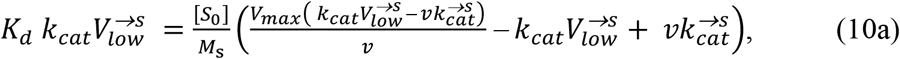

At this juncture, there is a need to clarify the relationship between 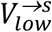 and *v*. They are the same from the first principle on account of which 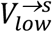 and *v* are directly proportional to [*S*_0_]/*M*_*s*_. Simplification of Eq. (10) gives the following:

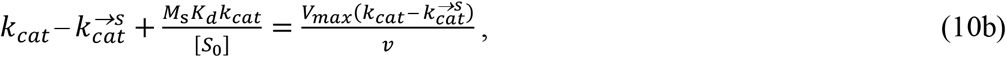

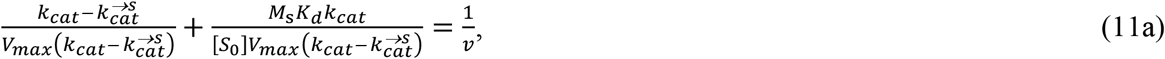

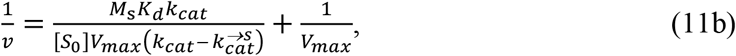

The slope (*S*_*lope*_) from the plot of 1/*v* versus 1/[*S*_0_] is given as follows:

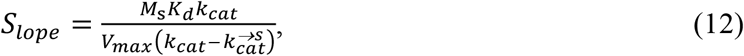

Meanwhile,

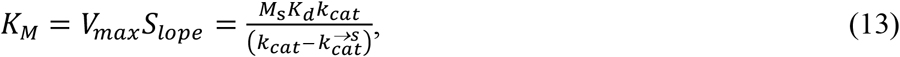

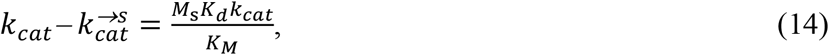

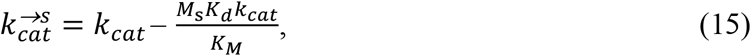

From Eq. (3):

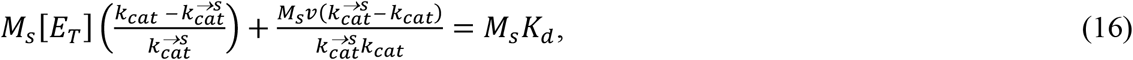

Let *M*_*s*_*K*_*d*_ = *K*_*dmc*_ such that, division of Eq. (16) by 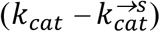 gives the following:

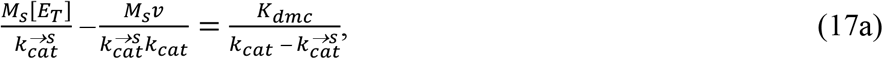

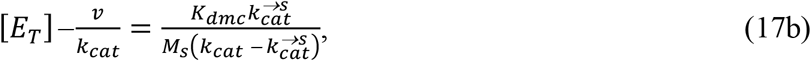

Note that [*E*_*F*_] = [*E*_*T*_]− *v*/*k*_*cat*_ and [*E*_*F*_] = *K*_*M*_[*ES*]/[*S*_*T*_] such that,

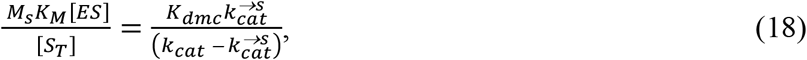

Returning [*ES*] to (*v*/*k*_*cat*_) in Eq. (18) and rearranging gives the following:

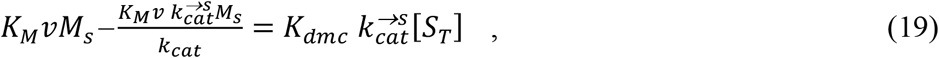

Solving for *K*_*dmc*_ gives the following:

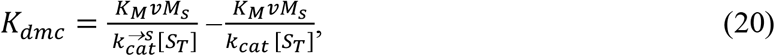

Meanwhile from Eq. (12) 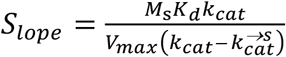. Thus,

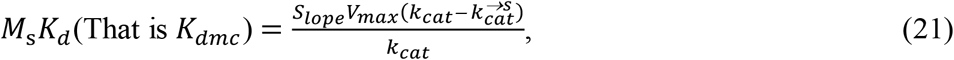

Therefore, Eqs (21) and (20) are the same. Hence,

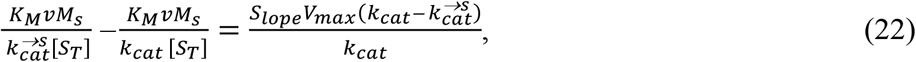

Expansion yields the following:

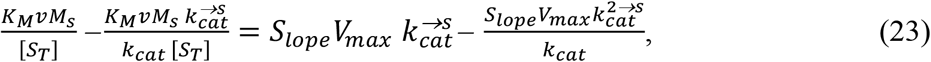

Rearrangement gives the following:

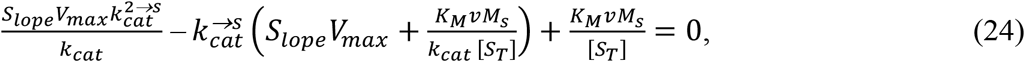

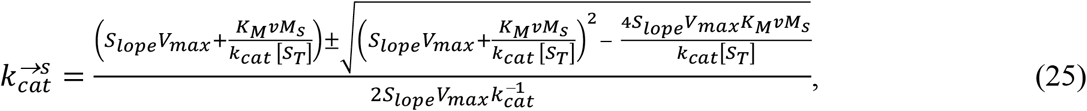

The possibility of two positive roots, α (the higher root) and β (the lower root) are derivable from Eq. (25). Preliminary investigation showed that 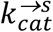 came out as *k*_*cat*_ represented as α and the former as ‘β’.

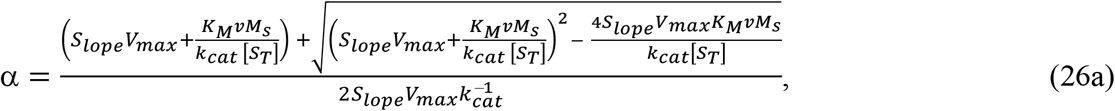

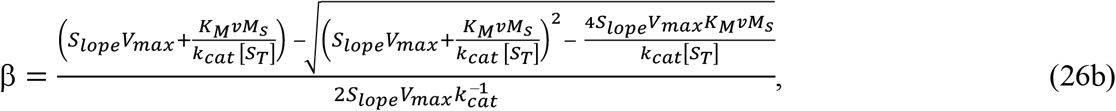

Note that after investigation alpha remains consistently, the zero-order (saturating) first-order catalytic rate constant, *k*_*cat*_ for the production of the product, while beta remains the pre-zero-order (unsaturating) first-order rate constant (Note: not the pseudo-first-order rate constant) for the utilization of the substrate. The pre-zero-order velocity, *v* of catalysis could be directly proportional to the molar concentrations of the enzyme for different molar concentration of the substrate up to such concentration by which the zero-order first-order rate constant is attained regardless of the highest concentration of the enzyme. Following the analysis, it is important to note that β remains the pre-zero-order (unsaturating) first-order rate constant for substrate utilization, and α remains the zero-order (saturating) first-order catalytic rate constant for product formation. Regardless of the maximum enzyme concentration, the pre-zero-order velocity of catalysis (v) may be directly proportional to the enzyme’s molar concentration when the substrate concentration falls within a range that permits reaching the zero-order first-order rate constant. To put it another way, the maximal enzyme concentration may be saturated by one of these amounts. Using a low enzyme concentration that can readily be saturated at uninhibiting substrate concentrations over the course of a brief assay period is advised by Udema (20xx).

### 2.1 Relating two different approaches for the determination of second-order rate constant for ES formation and first-order rate constant for its dissociation

#### 2.1.1 Second-order rate constant for ES formation for each [S_T_]

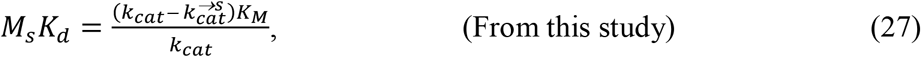

Recall that, the unit of *K*_*d*_ is molarity.

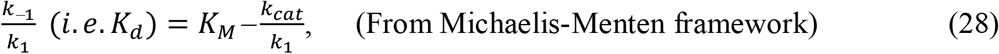

Thus,

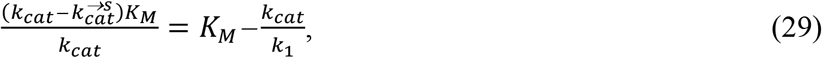

Solving for *k*_1_ yields the following:

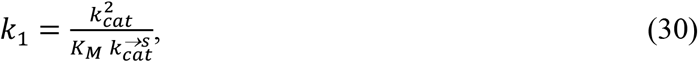

#### 2.1.2 First-order rate constant for ES dissociation for each [S_T_]

By equating *k*_−1_/*k*_1_with the right hand side of Eq. (27) one can solve for *k*_−1_. Thus,

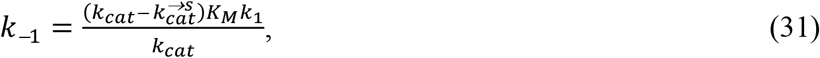

Substitution of Eq. (30) into Eq. (31) gives the following:

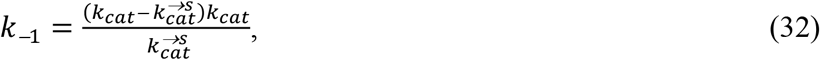

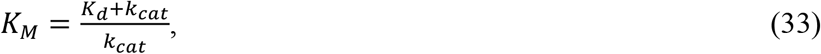

In the QSSA approximation, this constant (Eq. 33) determines the proportion of bound enzyme at a given substrate concentration (Berberan-Santos, 2010). But how? Does this imply that half of the enzyme is bound at this concentration? This conclusion is based on the definition of the Michaelis–Menten constant. The rate constant for the production of the product is not necessarily equivalent to the rate at which the enzyme attaches to the substrate and chemically converts it. The turnover rate (Berberan-Santos, 2010) is as follows:

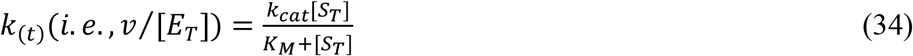

In accordance with Eq. (34), it is evident that *k*_(*t*)_ is a fraction of *k*_*cat*_. However, if almost the entire enzyme is present in bound form at saturation (where [*S*_*T*_] >[*E*_*T*_]), as defined, the former (*k*_*cat*_) is rarely less than one. This leads to the maximal turnover rate, *k*_*cat*_ which is also referred to as the turnover number. Additionally, turnover rates usually fall between 100 and 1,000 per second, with a maximum of about 10^6^/s (Berberan-Santos, 2010). The terms “turnover” and “numbers” are not confusing when rephrased in terms of magnitude as “turnover number.” This must, however, be in line with single-turnover kinetics, which forbids the catalytic cycling that is often discussed in the literature. Note that even if [*E*_*T*_]»[*S*_*T*_] and the latter are fully utilized within the chosen short assay duration, if the substrate’s molar mass is greater than the enzyme’s, the [*S*_*T*_] utilized per unit time per [*E*_*T*_] may be less than one ([*S*_*T*_]/[*E*_*T*_] ≡ *M*_2_/*M*_3_). This ratio may be greater than one if *M*_2_ is much greater than *M*_3_.

## 3. Experimental

### 3.1 Materials

#### 3.1.1 Chemicals

Human salivary alpha-amylase (EC 3.2.1.1) and insoluble potato starch were purchased from Sigma-Aldrich in the USA. Tris, 3,5-dinitrosalicylic acid, maltose, and sodium potassium tartrate tetrahydrate were purchased from Kem-Light Laboratories in Mumbai, India. Hydrochloric acid, sodium hydroxide, and sodium chloride were purchased from BDH Chemical Ltd. in Poole, England. Distilled water was purchased from a local market. The molar mass of the enzyme is 52kDa (Sugahara *et al*., 2013).

#### 3.1.2 Equipment

As in previous studies, an electronic weighing machine was purchased from Wenser Weighing Scale Limited, and a 721/722 visible spectrophotometer was purchased from Spectrum Instruments, China. A pH meter was purchased from Hanna Instruments, Italy.

### 3.2 Methods

The Bernfeld method (Bernfeld, 1955) was used to quantify the velocity of amylolysis given a a gelatinized substrate mass concentration ranging from 3 to 9 g/L. This theoretical and experimental research focuses solely on human salivary alpha-amylase. This study also, explored data from invertase (EC 3.2.1.26), a beta fructofuranosidase assay in the literature (Bowsk *et al*., 1971). Maltose was used as a standard at 540 nm with an extinction coefficient of 181 L/mol cm to assess the amount of reducing sugar produced by hydrolyzing the substrate at 310.15 K. It took three minutes to finish the experiment for each concentration of the substrate. In a Tris-HCl buffer solution at pH 6.6, the mass concentration of the human salivary alpha-amylase () was 3.24 mg/L.

### 3.3 Statistics

The enzyme was only used in duplicate tests. Because a minimum dataset of *n* ≥ 3 is required for nonparametric statistics and *n* ≥ 6 for parametric statistics, it was impossible to determine the standard deviation either parametrically or nonparametrically, much to the delight of advanced biostatisticians, since the mean of each duplicate was used for all calculations.

## 4. Results and discussion

### 4.1 Results

As shown in Table 1, the kinetic parameters (for human salivary alpha amylase, HSAA), including the ES dissociation constant (*K*_*d*(*mc*)_), the second-order rate constant, ***k***_***1***_ for ES formation, and the first-order rate constant for ES dissociation, increased with higher substrate concentrations. The first-order rate constant for substrate utilization, 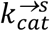, exhibited a dissimilar trend with higher [*S*_0_] (see Table 1). The zero-order equivalents of the other parameters increased with rising substrate concentrations, but the magnitude of the zero-order 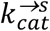 was the lowest. The magnitude of *K*_*d*(*mc*)_ increased as substrate concentration rose, indicating that an increase in product concentration diminished the enzyme’s affinity for the substrate.

**Table 1:** Kinetic constants in enzymatic catalysis by human salivary alpha amylase for each substrate concentration.

| $[S_0]/g/L$ | $K_{d(mc)}/g/L$ | $k_1/L/g.min$ | $k_{cat}^{\rightarrow s}/min$ | $k_{-1}/min$ |
| --- | --- | --- | --- | --- |
| 3 | 2.31113 | 5,452.4894 | 5,802.4687 | 1,2601.4121 |
| 4 | 2.57045 | 6,185.5497 | 5,114.8111 | 1,5899.6555 |
| 5 | 2.77473 | 6,918.2196 | 4,573.1301 | 1,9196.1568 |
| 6 | 2.93983 | 7,650.6880 | 4,135.3037 | 2,2491.7522 |
| 7 | 3.07606 | 8,383.0013 | 3,774.0595 | 2,5786.6169 |
| 8 | 3.19038 | 9,115.1381 | 3,470.9204 | 2,9080.7522 |
| 9 | 3.28764 | 9,846.8087 | 3,213.0124 | 3,2372.7577 |

As a reminder, $K_{d(mc)}$ is the ES dissociation constant measured in g/L (note L symbolizes the liter so as not to confuse 1, with lower case of L). Zero-order kinetic constants $k_1 = 15,082.9563$ L/g/min; $k_{-1} = 55,931.7566$ /min; $K_{d(mc)} = 3.7083$ g/L; $k_{cat} = 11,930.9885$ /min; Michaelis-Menten constant: 4.4993 g/L

All of the computed experimental parameters are displayed in Table 2, and they demonstrate an increasing trend as substrate concentrations rise. The trend of these parameters is similar to that of human alpha amylase. However, outliers in the invertase plots disrupt the trend according to the first or second higher substrate concentration reported in the literature and examined in this study, making comparisons challenging. In contrast to HSAA, the invertase and substrate concentrations were abnormally high. Despite the sucrose content being several times greater, the enzyme mass concentration of 5 g/L (100 mg/20 mL) was excessive. The exceptionally low 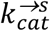 values were caused by the very high *k*_−1_ and *K*_*d*(*mc*)_ values, which were comparable to *K*_*M*_ This suggests that, in addition to the impact of the high sucrose concentration on viscosity, the presence of a high product concentration reduced the enzyme’s affinity for sucrose (Bowski *et al*., 1971).

**Table 2:**
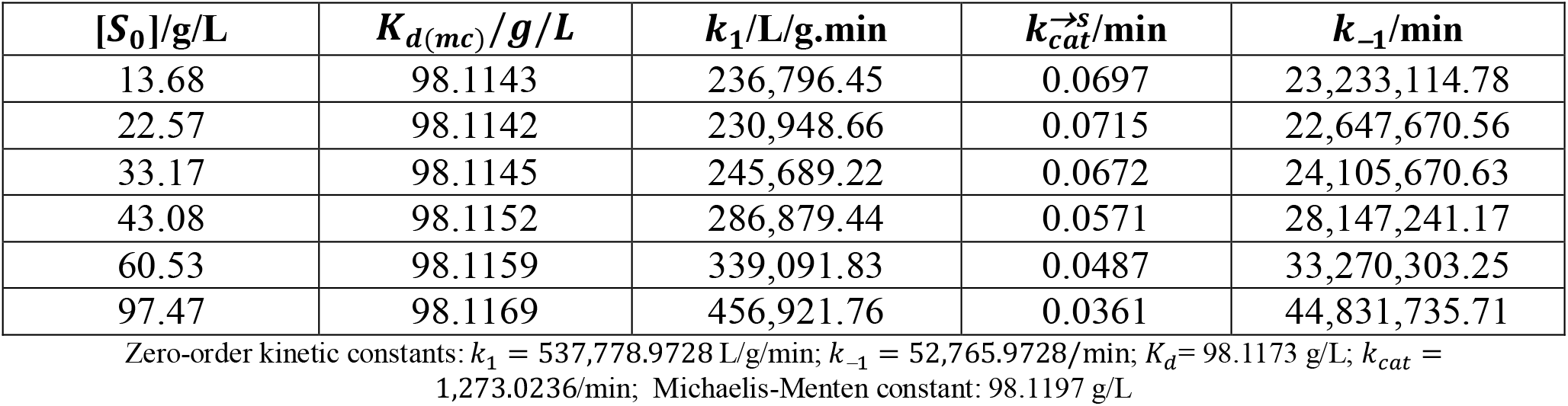
Kinetic constants in enzymatic catalysis by invertase (from baker’s yeast) for each substrate concentration Table1: Kinetic constants in enzymatic catalysis for each substrate concentration.

### 4.1 Cases in the literature where single turnover is utilized in mechanistic investigations *vis-à-vis* this Michaelian study

In connection to the results of this extremely detailed Michaelian test, it is important to consider cases in the literature where single turnover is utilized in mechanistic investigations of different types of enzymes under different conditions. Several pieces of literature record the belief that the chemistry of an enzymatic action is best observed on the basis of single-turnover kinetics. Examples are as follows: The single-turnover approach has been used in investigative experiments, as documented. These studies have revealed that the intrinsic rate at which MutY removes adenine from an OG:A substrate is at least six times faster than from a comparable G:A substrate. However, when [MutY] is much smaller than [DNA], OG:A substrates are not quantitatively converted to the final product due to inefficient turnover caused by slow product release. *Yet, the catalytic rate constant for the product is regarded as the turnover rate instead of the rate constant for the utilization (or conversion) of the substrate!* In contrast, MutY dissociates more easily from the product when the substrate is G:A, allowing for complete conversion under similar conditions. Kinetic results demonstrate significant differences in the glycosylase reaction catalyzed by MutY, *depending on the substrate’s characteristics* (Porello *et al*., 1998). One of such characteristics of interest is the degree of polymerization or rather the number of vulnerable site per substrate to be attacked by the enzymes. According to Martinelli *et al*. (2023), another example of a study in which the single-turnover concept is applied gave results that suggest that MtInhA displays a typical symmetry model (Monod *et al*., 1965) with free enzymes in equilibrium between two conformers, followed by NADH binding to one conformer and conformational change in the binary complex (Martinelli *et al*., 2023).

Single turnover kinetics of tryptophan hydroxylase: Evidence for a new intermediate in the reaction of the aromatic amino acid hydroxylases (Pavón *et al*., 2010). How does single turnover relate to catalytic rate constants? According to chemical quench analyses, the formation of 5-hydroxytryptophan occurs at a rate constant of 1.3/s. The total turnover is likely to be constrained by a subsequent slow step involving the release of either hydroxypterin or the hydroxylated amino acid, with a rate constant of 0.2/s (Pavón *et al*., 2010). The question then becomes which of these rate constants can define the turnover number while taking into account the requirement for a cosubstrate or cofactor. The two main products are Quinonoid Dihydrobiopterin (q-BH_2_) and 5-hydroxytryptophan (5-HTP). The main substrate remains tryptophan. If concern is expressed about a single turnover rate, neither 1.3 nor 0.2 typifies it. There is no clear statement tied exclusively to the turnover rate or number.

Therefore, it is reasonable to argue that the rate constant for the formation and release of the product, also known as the catalytic rate constant or the turnover number, is not suitable. Rather, the rate constant for substrate utilization should be more appropriate. It is impossible to refer to a single turnover if a substrate has many vulnerable bonds. Dipeptides, oligopeptides, polypeptides, *etc*., as well as disaccharides, trisaccharides, polysaccharides, *etc*., serve as instructive examples here. For further insight on this issue, one can consult the literature (Udema, 2026). Apart from thermodynamic issues related to the enzyme’s physical properties (Udema, 2016), one should realize that the molar masses of the enzyme, substrate, and product affect the computation of the catalytic rate constant, given the following equations: 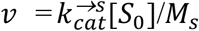 and 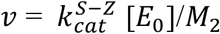 (is the sub-zero-order catalytic rate constant), 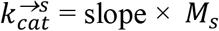 and 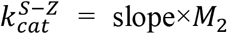. These equations supplement *V*_*Max*_*M*_2_/[*E*_0_]. Therefore, 1.3/s is approximately equivalent to 1/s. However, the terminology in the literature regarding the number of substrates makes it unclear how such translates to a single turnover rate. While the molar concentration of glucose produced by hydrolyzing maltose is twice that of maltose itself, the same cannot be said for any protein or polysaccharide. Such is particularly true for 1/s and 1.3/s.

The molar mass of a tryptophan hydrolase subunit is 51 kDa, depending on its location in the body (Sakowski *et al*., 2006). In contrast, the active tetrameric molar mass may be roughly four times greater. Without a doubt, the molar concentration of the new isomer is equal to the concentration of the isomerized reactant species in any enzyme-catalyzed isomerization reaction. Likewise, the turnover number is equal to the concentration of reactant species utilized (or transformed) per unit time, divided by the concentration of the enzyme. Consequently, the substrate turnover rate is equivalent to the catalytic rate constant for product creation only when the mole ratio of the enzymatic action product to one of the substrates is 1:1. The latter is a possibility in a perfect condition, in which generated experimental data are flawless. Additionally, the diffusion or translation of reacting species and their binding to one another occur over a different time frame than the chemical processes accelerated by the enzyme’s active site. Furthermore, the significant presence of the enzyme at concentrations several times higher than [*S*_0_] is consistent with the pre-steady-state or sub-*K*_*M*_ scenario. This implies that some enzyme molecules do not contribute to forming the complex that establishes the basis for related chemical and mechanistic questions. In contrast, not all substrate molecules form an enzyme-substrate complex in a zero-order scenario where substrate concentrations are much higher than [*K*_*M*_] and [*E*_*T*_]. Then does it mean that excess enzyme molecules don’t interfere with mechanistic studies?

### 4.2 Equilibrium issues

**I**n a study on Henri-Michaelis-Menten enzyme kinetics, the equilibrium notation is based on the equilibrium chemistry assumption. To obtain the enzyme–substrate complex (ECA) formulation, the researcher makes the following assumptions (Berberan-Santos, 2010):

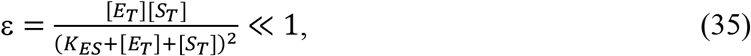

The ECA formulation of ES is obtained as (Berberan-Santos, 2010):

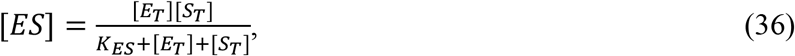

However, it is unclear whether Eq. (36) represents the equilibrium concentration of the enzyme-substrate complex (ES). This is in light of the conflicting claims about whether equilibrium kinetics is valid in the formulation of the Michaelian equation. When the enzyme-reactant intermediate concentration approaches the ultimate equilibrium concentration, a steady state is reached. The reactant with the lower binding constant is the first to do this. As a result, kinetic analyses that employ the Haldane relationship and the *k*_*cat*_/*K*_*M*_ ratio as specificity metrics are typically unreliable (Barnsley, 2022). Nevertheless, this work suggests that reactions catalyzed by enzymes are not ionic reactions involving distinct reactants, like AgNO_3_*(aq)* and HCl*(aq)*. Without a catalyst, these diffusion-driven reactions accelerate as the temperature rises. Conversely, after encounter-complex formation, enzyme-catalyzed events take center stage. The encounter-complex formation is diffusion-driven and may not always result in stable ES development. The second phase involves complex mechanistic chemistry, distortion of molecular orbitals, conformational and configurational reorientation, and bond formation and cleavage before product release. This phase may or may not take longer than the initial formation of the encounter complex. Some encounters and even unstable ES may be terminated during this period, briefly putting the system in a dynamic state. This influenced the study’s derivation, which started with Eq. (1).

Briggs-Haldane’s system is open from a thermodynamic and biological perspective, but MM kinetics is a closed system that causes the substrate and total reaction rates (*v*) to follow a hyperbolic function when the values are 18, 23, and 34. According to the mass action law, such behavior can only be anticipated if the substrate is in excess (Ariyawansha *et al*., 2018).

To create enzyme-substrate complexes that adhere to the mass action rule, the researchers devised a reliable plan involving the cyclic generation of reactants and enzymes. This approach effectively applies Michaelis–Menten kinetics to all biochemical processes involving single parameters. This formalism is a vital tool for solving several challenging environmental and healthcare issues. When developing the formalism, they employed rate and time perspectives to define the substrate concentration as [Product]^¾^ and the reaction rate (Ariyawansha *et al*., 2018). It is an essential tool to overcome some challenging healthcare and environmental issues (Ariyawansha *et al*., 2018). The interest in these pieces of information lies in the term “single parameter” and the question of how a fractional unit bearing [Product]^¾^ fits into the Michaelis– Menten framework.

### 4.3 The lowest and highest substrate concentration at which the rate constant for substrate utilization and production formation are equal

Two nonlinear regression equations of the polynomial type are obtained by plotting the proportion of the zero-order rate constant for product production and release. This is calculated as the ratio of *v*/[*E*_*T*_] to *k*_*cat*_. Then, the beta coefficient in Eq. (25) is plotted against [*S*_0_]. The equations are as follows:

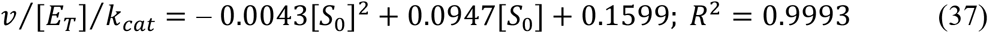

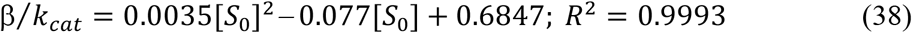

Relating equations (37) and (38) to each other yields another quadratic equation. This new equation has roots that correspond to the higher and lower substrate concentrations that produce the same results when substituted into the original equations. The other equation is given as follows:

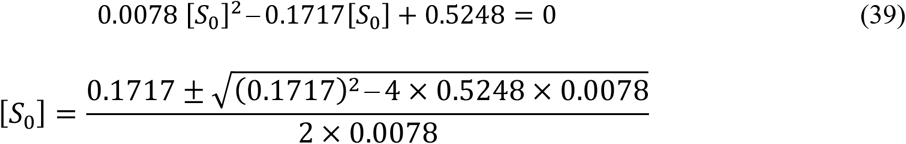

α = 18.34528043 g/L (The higher value of [*S*_0_])

β = 3.667540128 g/L (The lower value of [*S*_0_])

When the alpha and beta values are substituted, equations (37) and (38) provide 0.450036006 × *k*_*cat*_ and 0.449377387 × *k*_*cat*_, respectively. Either value supports the notion that there is a substrate concentration at which the rate constants for substrate utilization and product generation are equal. While the alpha value of the substrate concentration (18.34528043 g/L) is theoretically sound and supported by initial experimental findings, a significantly higher concentration of HSAA (approximately 0.00347826 g/L) would be required for the single-turnover assay. The substrate concentration’s beta value (3.667540128 g/L) falls within the range of substrate concentrations examined in this investigation (3 to 9 g/L). Its molar concentration is 5.682584642 M, which is comparable to the enzyme’s 5.610096774×*exp*.(−8) M. The concentration of the latter would need to be significantly higher than 0.00347826 g/L for the single-turnover assay. To fully convert all of the polymers to product with little to no cycling over the brief timeframe of the chosen assay, perhaps a concentration 100 times greater would be required.

## 5. Conclusion

Unlike other kinetic constants, the rate constant 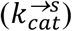 for substrate utilization decreases as substrate concentration increases: This is consistent with the long-standing observation that direct linearity shifts toward zero-order or hyperbolic Michaelian kinetics at higher substrate concentrations. However, there are substrate concentrations at which 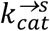 and *k*_*cat*_ are equal. The lower concentration falls within the range explored by the assay in this study and is preferable for higher enzyme concentrations, as it ensures single-turnover kinetics. For any future assays involving different enzymes, it is recommended that Michaelian and sub-Michaelian kinetic experiments be conducted and the results of cognate kinetic parameters be reported for comparison to aid the determination of the exact type of kinetic parameters.

## 6. Author Contribution

The sole author designed, analyzed, interpreted and prepared the manuscript.

## 7. Dedication

This study is dedicated to a father, a co-teacher, and his handsome baby boy. The father approached me in the third week of the month for help transporting his son to a clinic some kilometers away. Unfortunately, I was unable to help him during the first structural adjustment program in the early 1990s, when there was no industrial foundation or export-oriented economy. The co-teacher’s announcement the following day about his son’s death was devastating to me. The World Bank and the IMF need to adopt values that benefit everyone.

## 8. Disclaimer (Artificial Intelligence)

Author(s) hereby declare that no generative AI technologies such as Large Language Models (ChatGPT, COPILOT, *etc*.) and text-to-image generators have been used during the writing or editing of this manuscript.

## 9. Competing Interests

There are no known competing financial interests, non-financial interests, or personal ties that could have influenced the work presented in this article. The only issue is the monthly pension, which is much less than two USD per day and may not have been recommended by the World Bank or the IMF.

## 10. Acknowledgement

I am truly grateful to my siblings for their financial and in-kind support.

## Notes

### Competing Interest Statement

The authors have declared no competing interest.

## References

Adediran SA, Pratt RF. Some differences in turnover kinetics of penams and cephems catalyzed by classes A and C β-lactamases Proc. Nigerian Acad. Sci. 2022; 15 (1); 135, Doi: 10.57046/WOLL8646

Ariyawansha RTK, Basnayake BFA, Karunarathna AK, Mowjood MIM. Extensions to Michaelis-Menten Kinetics for Single Parameters. Sci Rep. 2018; 8(1): 16586, Doi: 10.1038/s41598-018-34675-2.

Banwait JK, Islam L, Lucius AL. Single turnover transient state kinetics reveals processive protein unfolding catalyzed by Escherichia coli ClpB eLife 2024; 13: RP99052, Doi: 10.7554/eLife.99052.2

Barnsley EA. Henri-Michaelis-Menten kinetics of reversible enzymatic reactions, and the determination of rate constants from kinetic constants Sci Prog. 2022; 105(2): 368504221100027, Doi: 10.1177/00368504221100027.

Berberan-Santos MN. A general treatment of Henri-Michaelis-Menten enzyme kinetics: Exact series solution and approximate analytical solutions *MATCH Commun*. Math. Comput. Chem. 2010; 63: 283–318, DOI: 10.48550/arXiv.0905.0874

Bergman WN, Sousa MC. A fluorescent reporter and single-turnover kinetics reveal insight into BAM complex function Proc. Natl. Acad. Sci. U.S.A. 2025; 122 (52): e2514687122, Doi: 10.1073/pnas.2514687122.

Bernfeld P. Amylases, alpha and beta Methods Enzymology. 1959; 1:149–158. Doi: 10.1016/00766879(55)01021-5

Bowski L, Saini R, Ryu DY, Vieth WR. Kinetic modeling of the hydrolysis of sucrose by invertase Biotechnol Bioeng. 1971;13(5):641–656. Doi: 10.1002/bit.260130505.

Dangerfield TL, Zhou J, Neumeyer J, Eickbush T, Johnson KA. Single-turnover kinetic analysis of non-long terminal repeat retrotransposition defines the pathway and rate constants leading to second-strand synthesis, Nucleic Acids Res 2026; 54 (6): 223, Doi: 10.1093/nar/gkag223

Kanazhevskaya LY, Alekseeva IV, Fedorova OS. A single-turnover kinetic study of DNA demethylation catalyzed by Fe(II)/α-Ketoglutarate-dependent dioxygenase AlkB. Molecules. 2019; 24(24):4576, Doi: 10.3390/molecules24244576

Martinelli LKB, Rotta M, Bizarro CV, Machado P, Basso LA. Single turnover of transient of reactants supports a complex interplay of conformational states in the mode of action of *Mycobacterium tuberculosis* enoyl reductase. Future Pharmacol. 2023; 3(2):379–391. Doi: 10.3390/futurepharmacol3020023

Monod J, Wyman J, Changeux JP. On the nature of allosteric transitions: A plausible model. 1965; J. Mol. Biol. 12: 88–118, Doi: 10.1016/S0022-2836(65)80285-6

Pavon JA, Eser B, Huynh MT, Fitzpatrick PF. Single turnover kinetics of tryptophan hydroxylase: Evidence for a new intermediate in the reaction of the aromatic amino acid hydroxylases. Biochemistry 2010; 49(35):7563–71. Doi: 10.1021/bi100744r.

Porello SL, Leyes AE, David SS. Single-turnover and pre-steady-state kinetics of the reaction of the adenine glycosylase MutY with mismatch-containing DNA substrates Biochemistry 1998; 37(42): 14756–14764, Doi: 10.1021/bi981594+

Sakowski SA, Geddes TJ, Thomas DM, Levi E, Hatfield JS, Kuhn DM. Differential tissue distribution of tryptophan hydroxylase isoforms 1 and 2 as revealed with monospecific antibodies. Brain Res. 2006; 1085(1): 11–18. Doi: 10.1016/j.brainres.2006.02.047.

Sugahara M, Takehira M, Yutani K. Effect of heavy atoms on the thermal stability of alphaamylase from *Aspergillus oryzae*, PLoS One. 2013; 2(2013):1–7. Doi: 10.1371/journal.pone.0057432

Tang JY. On the relationships between Michaelis–Menten kinetics, reverse Michaelis–Menten kinetics, equilibrium chemistry approximation kinetics and quadratic kinetics *Geosci*. Model Dev. Discuss 2015; 8: 7663–7691, Doi: 10.5194/gmdd-8-7663-2015

Udema II. Lineweaver Burk plot, rate constant and mathematical relationship between molar mass and free energy of activation: “To be or not to be.” J. Sci. Res. Rep. 2016; 10: 1–8. Doi: 10.9734/JSRR/2016/24857

Udema II. Turnover and Catalytic Cycle Frequency Determination Based on Molar Mass-Dependent Model Equations Int. J. Adv. Multidisc. Res. Stud. 2026; 6(2):1597–1607, Doi: 10.62225/2583049X.2026.6.2.6142

